# Development of Motif-Specific Monoclonal Antibodies for Global Protein Citrullination Detection with Minimal Cross-Reactivity to Homocitrullination

**DOI:** 10.1101/2025.03.27.645732

**Authors:** Sophia Laposchan, Erik Riedel, Andrew Flatley, Regina Feederle, Chien-Yun Lee

## Abstract

Protein citrullination, a post-translational modification (PTM) catalyzed by peptidylarginine deiminases (PADs), plays critical roles in biological processes such as immunity, gene regulation, and inflammation. Dysregulated citrullination is implicated in diseases including rheumatoid arthritis, multiple sclerosis, and cancer, making it a potential biomarker and therapeutic target. Immunodetection is the most commonly used technique to study citrullination. However, most commercially available antibodies against citrullination are either protein-specific or lack the sensitivity and specificity of the broad variety of modified proteins. In addition, existing anti-pan citrullination antibodies often fail to distinguish citrullination from homocitrullination, a chemically similar modification on lysine. This cross-reactivity limits their utility in deciphering the distinct biological roles of these PTMs. To address these challenges, we employed a motif-based strategy to generate monoclonal antibodies against citrullination. This approach leverages PAD enzyme sequence preferences from human proteome, enhancing specificity for citrullination while minimizing cross-reactivity with homocitrullination. We immunized rats with a pool of over 490,000 citrullinated peptides, designed to represent common citrullination motifs in human tissue proteomes. Two monoclonal antibody clones were established and validated for sensitivity and specificity using ELISA and western blot against *in vitro* citrullinated and homocitrullinated proteomes, as well as ionomycin-activated human neutrophils. Both clones demonstrated great sensitivity to diverse citrullinated proteins, robust discrimination against homocitrullination, and quantitative readout in biological samples. These novel antibodies provide powerful tools for studying global citrullination dynamics and hold promise for biomarker discovery and diagnostic applications in diseases involving PAD dysregulation.

## Introduction

Protein citrullination is a post-translational modification (PTM) catalyzed by peptidylarginine deiminases (PADs) in the presence of calcium, converting arginine residues into citrulline. Its dynamics are critical in diverse biological processes, including innate immunity^1^, gene regulation^2^, inflammation^3^, and apoptosis^4^. Dysregulation of citrullination is implicated in various diseases, such as rheumatoid arthritis (RA), multiple sclerosis, and cancer^5,6^. Detecting citrullination is critical for understanding its cellular functions and the activities of PAD enzymes in both health and disease. This knowledge is essential for exploring its potential as a biomarker and therapeutic target. However, current methods for globally detecting citrullination remain limited.

Mass spectrometry (MS) and immunodetection are the primary approaches for PTM analysis, but both face unique challenges in detecting citrullination^7^. MS, while offering site-specific and quantitative data, is hindered by the small mass change of citrullination (0.98 Da) and the identical mass of deamidation on Gln and Asn. This increases false positives, especially when incorrect monoisotopic precursors are selected or incorrect localization of the 0.98 Da modification occurs on peptides containing Arg, Gln, or Asn^8,9^. Although advanced computational strategies^8,10,11^ have been proposed to improve data analysis, MS detection requires specialized instrumentation and expertise in sample preparation and data interpretation, limiting its general accessibility.

Compared to MS, immunodetection is widely used in almost every laboratory due to its accessibility and ease of use. While several anti-citrullination antibodies exist, most are limited to detecting citrullination sites on specific proteins, such as histones and myelin basic protein (MBP) (**Table 1**). Detecting global changes in citrullination is crucial for understanding PAD enzyme activity and the broader functional roles of citrullination. However, developing sensitive and specific anti-pan citrullination antibodies is challenging due to the subtle structural difference between citrulline and arginine^7^. The choice of immunogen is therefore critical for generating anti-pan citrullination antibodies that can detect small changes in citrullination without being restricted to specific proteins.

**Table 1.**
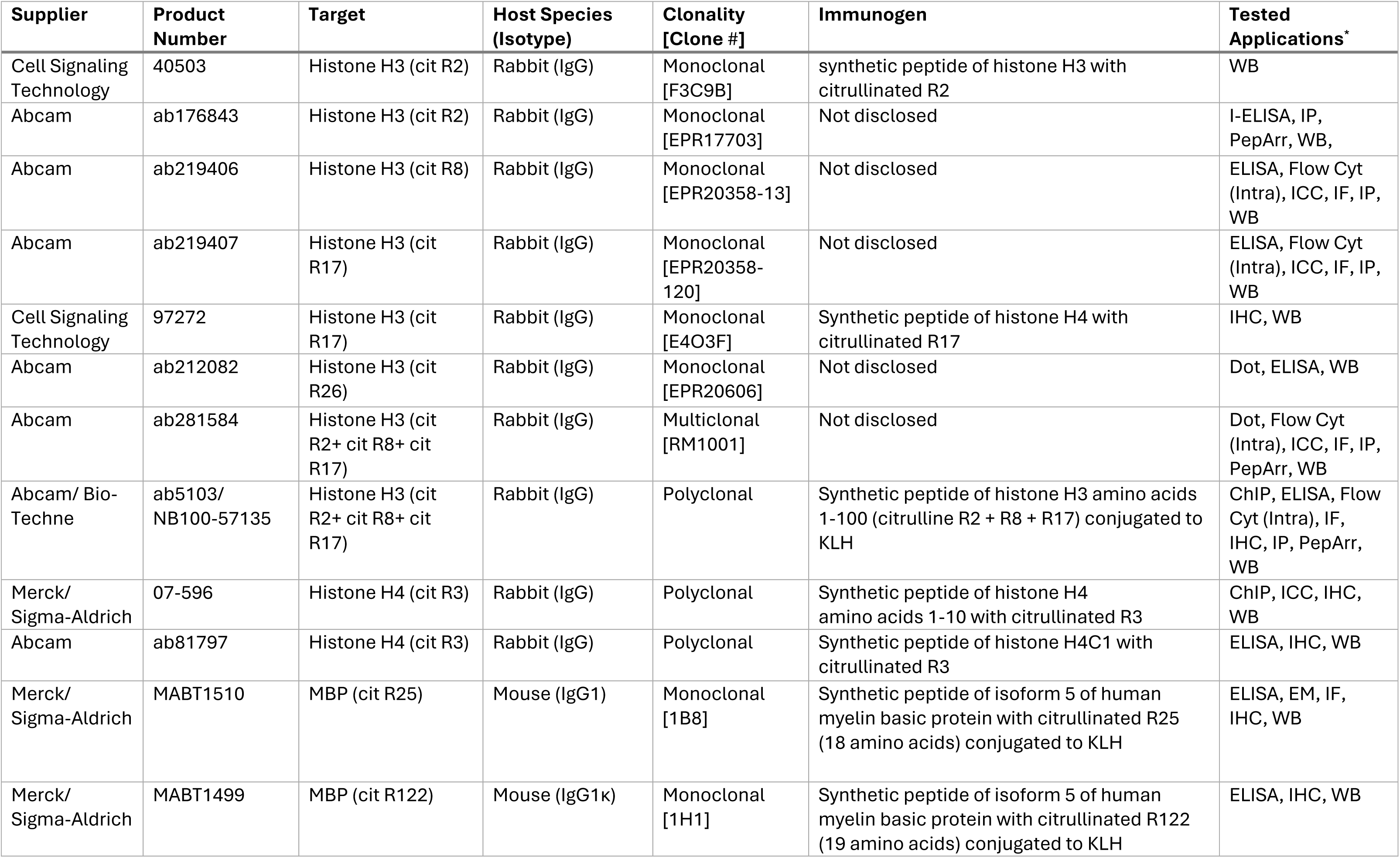
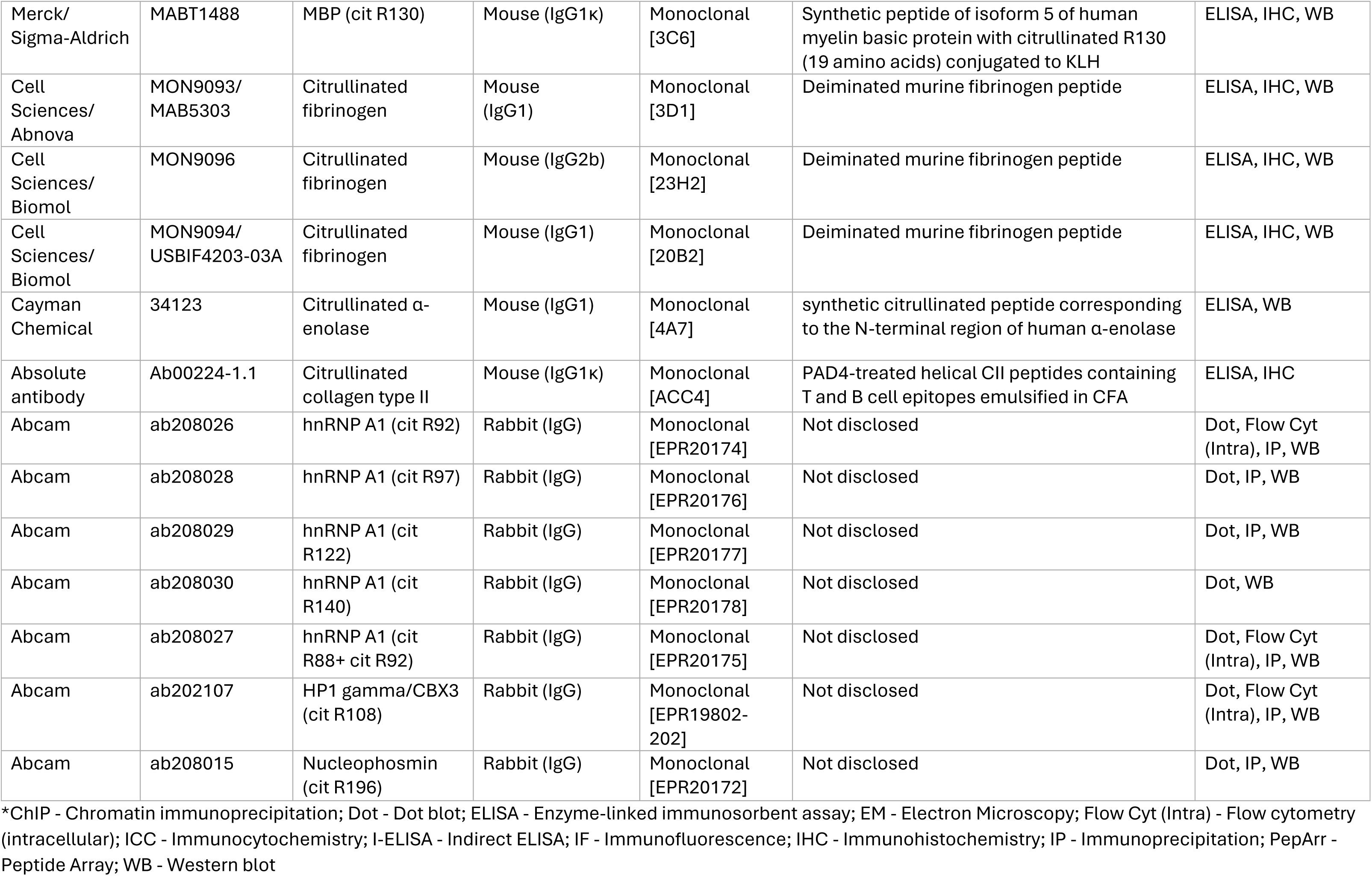
Summary of commercially available protein-specific anti-citrullination antibodies.

| Supplier | Product Number | Target | Host Species (Isotype) | Clonality [Clone #] | Immunogen | Tested Applications* |
| --- | --- | --- | --- | --- | --- | --- |
| Cell Signaling Technology | 40503 | Histone H3 (cit R2) | Rabbit (IgG) | Monoclonal [F3C9B] | synthetic peptide of histone H3 with citrullinated R2 | WB |
| Abcam | ab176843 | Histone H3 (cit R2) | Rabbit (IgG) | Monoclonal [EPR17703] | Not disclosed | I-ELISA, IP, PepArr, WB, |
| Abcam | ab219406 | Histone H3 (cit R8) | Rabbit (IgG) | Monoclonal [EPR20358-13] | Not disclosed | ELISA, Flow Cyt (Intra), ICC, IF, IP, WB |
| Abcam | ab219407 | Histone H3 (cit R17) | Rabbit (IgG) | Monoclonal [EPR20358-120] | Not disclosed | ELISA, Flow Cyt (Intra), ICC, IF, IP, WB |
| Cell Signaling Technology | 97272 | Histone H3 (cit R17) | Rabbit (IgG) | Monoclonal [E4O3F] | Synthetic peptide of histone H4 with citrullinated R17 | IHC, WB |
| Abcam | ab212082 | Histone H3 (cit R26) | Rabbit (IgG) | Monoclonal [EPR20606] | Not disclosed | Dot, ELISA, WB |
| Abcam | ab281584 | Histone H3 (cit R2+ cit R8+ cit R17) | Rabbit (IgG) | Multiclonal [RM1001] | Not disclosed | Dot, Flow Cyt (Intra), ICC, IF, IP, PepArr, WB |
| Abcam/ Bio-Techne | ab5103/ NB100-57135 | Histone H3 (cit R2+ cit R8+ cit R17) | Rabbit (IgG) | Polyclonal | Synthetic peptide of histone H3 amino acids 1-100 (citrulline R2 + R8 + R17) conjugated to KLH | ChIP, ELISA, Flow Cyt (Intra), IF, IHC, IP, PepArr, WB |
| Merck/ Sigma-Aldrich | 07-596 | Histone H4 (cit R3) | Rabbit (IgG) | Polyclonal | Synthetic peptide of histone H4 amino acids 1-10 with citrullinated R3 | ChIP, ICC, IHC, WB |
| Abcam | ab81797 | Histone H4 (cit R3) | Rabbit (IgG) | Polyclonal | Synthetic peptide of histone H4C1 with citrullinated R3 | ELISA, IHC, WB |
| Merck/ Sigma-Aldrich | MABT1510 | MBP (cit R25) | Mouse (IgG1) | Monoclonal [1B8] | Synthetic peptide of isoform 5 of human myelin basic protein with citrullinated R25 (18 amino acids) conjugated to KLH | ELISA, EM, IF, IHC, WB |
| Merck/ Sigma-Aldrich | MABT1499 | MBP (cit R122) | Mouse (IgG1κ) | Monoclonal [1H1] | Synthetic peptide of isoform 5 of human myelin basic protein with citrullinated R122 (19 amino acids) conjugated to KLH | ELISA, IHC, WB |
| Merck/<br>Sigma-Aldrich | MABT1488 | MBP (cit R130) | Mouse (IgG1κ) | Monoclonal<br>[3C6] | Synthetic peptide of isoform 5 of human myelin basic protein with citrullinated R130 (19 amino acids) conjugated to KLH | ELISA, IHC, WB |
| Cell<br>Sciences/<br>Abnova | MON9093/<br>MAB5303 | Citrullinated<br>fibrinogen | Mouse<br>(IgG1) | Monoclonal<br>[3D1] | Deiminated murine fibrinogen peptide | ELISA, IHC, WB |
| Cell<br>Sciences/<br>Biomol | MON9096 | Citrullinated<br>fibrinogen | Mouse (IgG2b) | Monoclonal<br>[23H2] | Deiminated murine fibrinogen peptide | ELISA, IHC, WB |
| Cell<br>Sciences/<br>Biomol | MON9094/<br>USBIF4203-03A | Citrullinated<br>fibrinogen | Mouse (IgG1) | Monoclonal<br>[20B2] | Deiminated murine fibrinogen peptide | ELISA, IHC, WB |
| Cayman<br>Chemical | 34123 | Citrullinated α-enolase | Mouse (IgG1) | Monoclonal<br>[4A7] | synthetic citrullinated peptide corresponding to the N-terminal region of human α-enolase | ELISA, WB |
| Absolute<br>antibody | Ab00224-1.1 | Citrullinated<br>collagen type II | Mouse (IgG1κ) | Monoclonal<br>[ACC4] | PAD4-treated helical CII peptides containing T and B cell epitopes emulsified in CFA | ELISA, IHC |
| Abcam | ab208026 | hnRNP A1 (cit R92) | Rabbit (IgG) | Monoclonal<br>[EPR20174] | Not disclosed | Dot, Flow Cyt<br>(Intra), IP, WB |
| Abcam | ab208028 | hnRNP A1 (cit R97) | Rabbit (IgG) | Monoclonal<br>[EPR20176] | Not disclosed | Dot, IP, WB |
| Abcam | ab208029 | hnRNP A1 (cit<br>R122) | Rabbit (IgG) | Monoclonal<br>[EPR20177] | Not disclosed | Dot, IP, WB |
| Abcam | ab208030 | hnRNP A1 (cit<br>R140) | Rabbit (IgG) | Monoclonal<br>[EPR20178] | Not disclosed | Dot, WB |
| Abcam | ab208027 | hnRNP A1 (cit<br>R88+ cit R92) | Rabbit (IgG) | Monoclonal<br>[EPR20175] | Not disclosed | Dot, Flow Cyt<br>(Intra), IP, WB |
| Abcam | ab202107 | HP1 gamma/CBX3<br>(cit R108) | Rabbit (IgG) | Monoclonal<br>[EPR19802-202] | Not disclosed | Dot, Flow Cyt<br>(Intra), IP, WB |
| Abcam | ab208015 | Nucleophosmin<br>(cit R196) | Rabbit (IgG) | Monoclonal<br>[EPR20172] | Not disclosed | Dot, IP, WB |

\*ChIP - Chromatin immunoprecipitation; Dot - Dot blot; ELISA - Enzyme-linked immunosorbent assay; EM - Electron Microscopy; Flow Cyt (Intra) - Flow cytometry (intracellular); ICC - Immunocytochemistry; I-ELISA - Indirect ELISA; IF - Immunofluorescence; IHC - Immunohistochemistry; IP - Immunoprecipitation; PepArr - Peptide Array; WB - Western blot

The earliest approach to address this challenge involves using chemically modified citrullinated peptides as immunogen to enhance immunogenicity, producing anti-modified citrulline antibodies (AMC)^12^. While highly sensitive, these antibodies require additional derivatization steps on the samples and are only applicable to western blotting and ELISA. Other strategies involve single citrullinated peptides or citrulline-conjugated carrier proteins (e.g., KLH or BSA), but these often suffer from sequence context-dependent recognition, limiting their ability to detect a broad range of citrullinated proteins. Another widely used antibody, F95, was therefore generated using a peptide with 10 citrullines as an immunogen and has demonstrated utility in detecting citrullination in human brain^13^. **Table 2** summarizes the immunogens and applications of commercially available anti-pan citrullination antibodies.

**Table 2.**
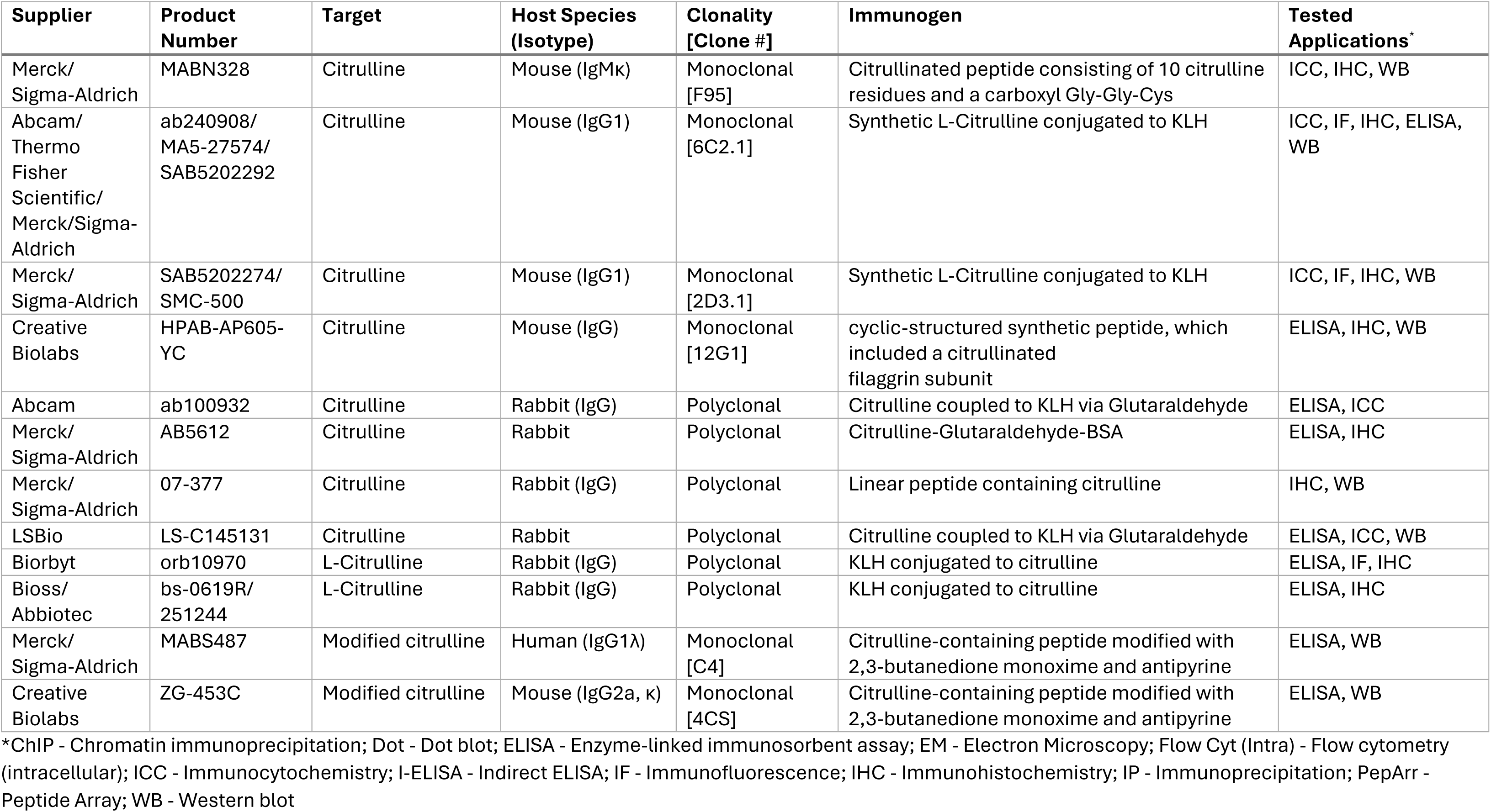
Summary of commercially available anti-pan citrullination antibodies.

Nevertheless, most anti-pan citrullination antibodies, including AMC and F95, often fail to distinguish between citrullination on Arg and homocitrullination on Lys due to the structural similarity of the modified residues^14,15^. Homocitrullination, also known as lysine carbamylation, occurs when lysine is chemically modified by cyanate, a potential byproduct of inflammation^16^. While PAD activity and cyanate levels are both elevated in inflammatory diseases such as RA, citrullination and homocitrullination are distinct modifications. Discriminating between these PTMs is essential for understanding their respective roles in disease mechanisms.

To address these challenges, we adopted a motif-based approach to generate anti-pan citrullination antibodies. Similar approaches have been successfully employed to create pan-PTM antibodies for phosphorylated S/T on the consensus motif of AGC kinase subfamilies (Akt and PKC)^17^, methylarginine GAR motif^18^, and ubiquitin remnant motif (K-ε-GG)^19^. A key distinction between citrullination and homocitrullination is that citrullination is catalyzed by PAD enzymes, which exhibit sequence-specific preferences adjacent to citrullination sites. Antibodies targeting PAD-specific motifs can potentially enhance specificity for citrullination while reducing cross-reactivity with homocitrullination. A similar concept has been used before in which polyclonal anti-citrullination antibodies recognizing the R(cit)GG motif were generated^20^. However, their sensitivity was limited because this motif accounted for only ∼20% of citrullinated substrates in their study.

Here, we developed monoclonal antibodies by immunizing with a pool of citrullinated peptides representing over 490,000 combinations to represent common citrullination motifs observed in human proteomes (**Figure 1A**). We established two stable clones, validated their sensitivity and specificity using direct ELISA as well as western blotting against *in vitro* citrullinated/ homocitrullinated proteomes, PAD isozyme-treated citrullinome, and ionomycin-activated neutrophils. Both clones demonstrated great sensitivity to various citrullinated proteins and specificity for citrullination over homocitrullination. These antibodies hold promise for advancing citrullination research and supporting biomarker discovery and diagnostics.

**Figure 1.**
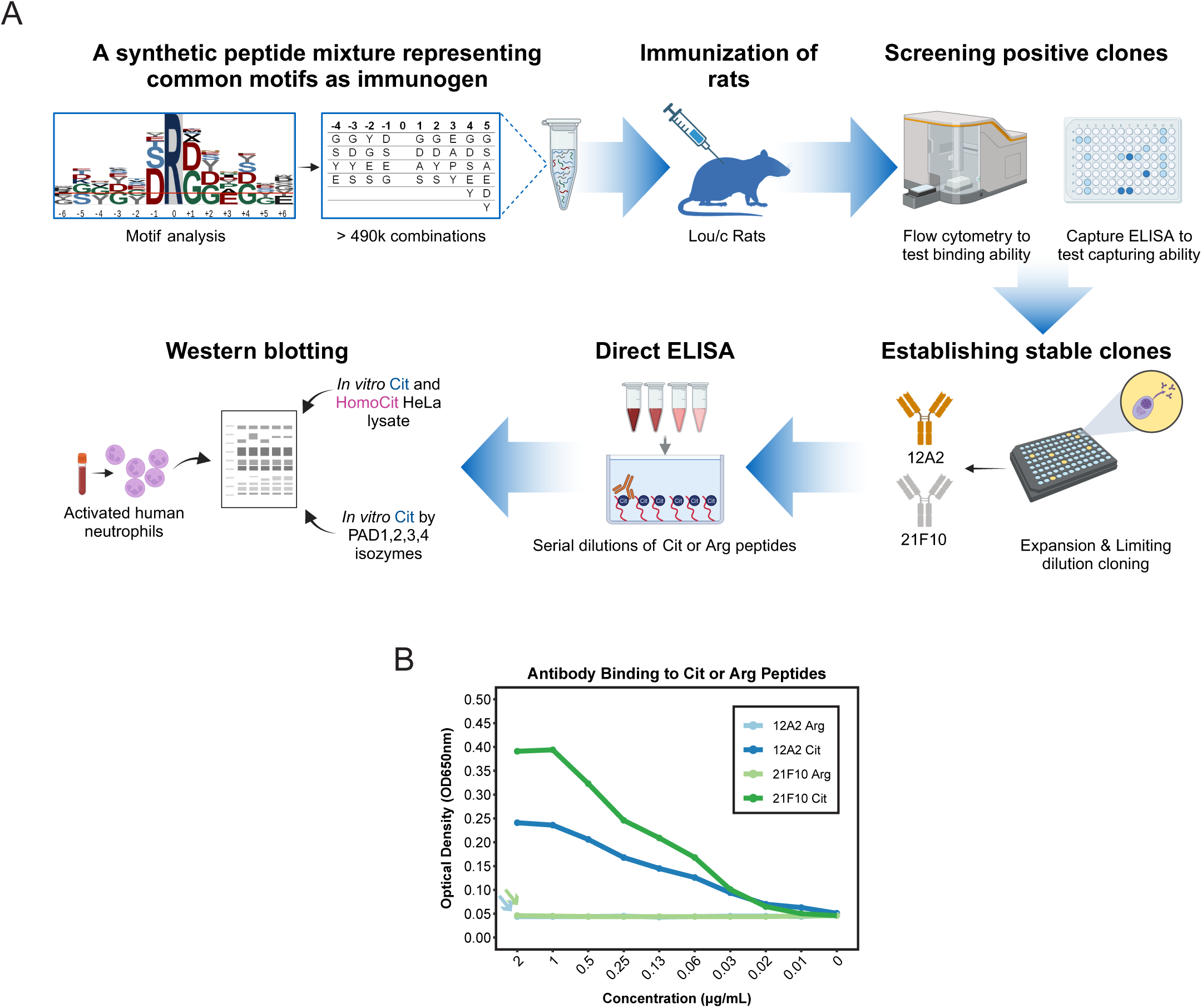
Generation of motif-specific monoclonal anti-pan citrullination antibodies. Schematic overview of the study design. Arg, arginine; Cit, citrulline/citrullination; HomoCit, homocitrulline/homocitrullination. (B) Binding specificity of custom monoclonal antibodies 12A2 and 21F10 toward a peptide mixture containing either citrulline (Cit) or arginine (Arg).

## Materials and Methods

### Generation of monoclonal antibodies against citrullinated peptides

A pool of 10aa-long peptides (xxxx-cit-xxxxx; x= mix of 4-6 different aa at each position) against citrullinated motifs identified from human proteome analyses (**Table 3**) were synthesized and coupled to ovalbumin (Ova) or biotin (Peps4LS, Heidelberg, Germany). Lou/c rats were immunized subcutaneously (s.c.) and intraperitoneally (i.p.) with a mixture of 40 µg Ova-coupled peptides in 400 µl PBS, 5 nmol CpG2006 (TIB MOLBIOL, Berlin, Germany), and 400 µl incomplete Freund’s adjuvant. Animals were boosted i.p. and s.c. with the same antigen preparation after 4-7 weeks, and a final boost without Freund’s adjuvant was given 13 weeks after primary immunization. Fusion of the myeloma cell line P3X63-Ag8.653 with the rat immune spleen cells was performed 3 days after the final boost using polyethylene glycol 1500 according to standard procedure^21^. After fusion, the cells were plated in 96-well plates using RPMI 1640 with 15% fetal calf serum, 1% penicillin/streptomycin, 1% glutamine, 1% pyruvate, 1x non-essential amino acids, 2% HCS (Capricorn) and HAT media supplement (Hybri-Max, Sigma-Aldrich). Hybridoma supernatants from 48 plates were screened 10 days later in a flow cytometry assay (iQue, Sartorius) on biotinylated citrullinated peptides captured on streptavidin beads (PolyAN, Berlin) and incubated for 90 min with hybridoma supernatant and Atto-488-coupled isotype-specific monoclonal mouse-anti-rat IgG secondary antibodies (TIB173 IgG2a, TIB174 IgG2b, TIB170 IgG1 all from ATCC, R-2c IgG2c homemade). A pool of biotinylated peptides containing arginine instead of citrulline was used for negative screening. Antibody binding was analyzed using ForeCyt software (Sartorius). Hybridoma cells of clones CIT 12A2 (rat IgG2c) and CIT 21F10 (rat IgG2b) were subcloned by limiting dilution to obtain stable monoclonal cell lines.

**Table 3.** Frequency table at each position of the pool of synthetic peptides.

| -4 |  | -3 |  | -2 |  | -1 |  | 0* | 1 |  | 2 |  | 3 |  | 4 |  | 5 |  |
| --- | --- | --- | --- | --- | --- | --- | --- | --- | --- | --- | --- | --- | --- | --- | --- | --- | --- | --- |
| G | 25% | G | 25% | G | 25% | D | 40% |  | G | 40% | G | 40% | E | 35% | G | 20% | G | 20% |
| S | 25% | D | 25% | S | 25% | S | 40% |  | D | 40% | D | 40% | A | 25% | D | 20% | D | 16% |
| Y | 25% | S | 25% | Y | 25% | G | 10% |  | S | 10% | Y | 10% | P | 25% | Y | 20% | S | 16% |
| E | 25% | Y | 25% | E | 25% | E | 10% |  | A | 10% | S | 10% | Y | 15% | S | 20% | Y | 16% |
|  |  |  |  |  |  |  |  |  |  |  |  |  |  |  | E | 20% | A | 16% |
|  |  |  |  |  |  |  |  |  |  |  |  |  |  |  |  |  | E | 16% |

\*Position of Citrulline or Arginine.

Animal experiments were conducted in accordance with the German animal welfare law and performed with permission and in accordance with all relevant guidelines and regulations of the district government of Upper Bavaria (Bavaria, Germany; Animal protocol number ROB-55.2Vet-2532.Vet_03-17-68).

### ELISA assay

Hybridoma supernatants of antibody clones 12A2 and 21F10 were tested in an enzyme-linked immunoassay (ELISA) for binding to biotinylated peptides containing citrulline or arginine as negative control. Serial dilutions of peptide (2ug/ml) in PBS with 20% fetal bovine serum (FBS) were added to avidin-coated plates overnight. After blocking with 2% FBS in PBS, hybridoma supernatants (1:10 dilution) were added for 30 min. After one wash with PBS, bound antibody was detected with HRP-coupled anti-rat kappa light chain antibody (TIB 172, ATCC). HRP was visualized with ready-to-use TMB substrate (1-StepTM Ultra TMB-ELISA, Thermo Fisher) and the absorbance was measured at 650nm with a microplate reader (Spark, Tecan).

### Cell culture

HeLa cells were cultivated in Dulbecco’s Modified Eagle Medium (DMEM) supplemented with 10% FBS. Prior to harvest, cells were washed twice with ice cold PBS and then lysed with a mild lysis buffer (50 mM Tris/HCl pH 7.5, 5% glycerol, 1.5 mM MgCl_2_, 150 mM NaCl, 1 mM Na_3_VO_4_, 25 mM NaF, 0.8% IGEPAL, 1 mM DTT). Cell lysate was cleared by ultracentrifugation for 1 h at 150,000 xg at 4 °C. Protein concentration of the lysate was determined using Pierce BCA Protein Assay Kit (Thermo Fisher Scientific, Rockford, IL, USA) according to the manufacturer’s instructions.

### *In vitro* protein homocitrullination assay

5 mg of HeLa lysate (5 μg/uL) was diluted with 1.5 mL PBS and 2.5 mL 2M potassium cyanate solution in PBS, resulting in 1M final concentration of potassium cyanate, and incubated overnight at 37 °C with shaking at 600 rpm. To remove residual cyanate before further analysis, buffer was exchanged using Amicon Ultra 10 kDA MWCO filters (Merck, Darmstadt, Germany). 5 mL of homocitrullinated lysate were filled up to 15 mL with PBS and centrifuged at 3224 xg until concentrated to ∼1 mL. Five iterations of centrifugation and dilution were performed to reduce the cyanate concentration to <0.01 mM. Protein concentration was determined again using Pierce BCA Protein Assay Kit. The resulting sample was then lyophilized and reconstituted to the desired concentration for further analysis. A control sample was prepared without potassium cyanate in the solution. For western blots, homocitrullinated samples were diluted five times with control sample to obtain a similar level of modification as the citrullinated samples.

### *In vitro* protein citrullination assay

5 mg of HeLa lysate (5 μg/uL) was diluted with 4 mL PAD buffer (79 mM Tris pH 7.6, 10 mM CaCl_2_, 2.5 mM DTT in final dilution) and incubated with recombinant PAD4 enzyme (kind gift from Prof. Hui-Chih Hung)^22^ at a 1:100 enzyme:protein ratio overnight at 37 °C with shaking at 600 rpm. For inactivation of PAD enzyme, samples were subsequently heated for 10 min at 80°C. Buffer exchange and lyophilization was carried out as described above.

For comparison of the citrullination patterns of different PAD isozymes, 200 μg of HeLa lysate (5 μg/μL) was diluted with 160 μL PAD buffer and incubated with recombinant PAD1 (Cay10784, Biomol), PAD3 (Cay10786, Biomol), PAD2, or PAD4 (both kindly provided by Prof. Hui-Chih Hung) enzyme at 1:100 enzyme:protein ratio overnight at 37 °C with shaking at 600 rpm. Enzymes were inactivated by heating samples for 10 min at 80°C. A control sample without any PAD was prepared analogously. Aliquots of the samples were lyophilized before western blot analysis.

### Neutrophils isolation and activation

Peripheral neutrophils were isolated from 25 mL of whole blood collected from a healthy volunteer using the EasySep Direct Human Neutrophil Isolation Kit (STEMCELL Technologies) via immunomagnetic negative selection, following the manufacturer’s protocol. After isolation, the cells were washed with HBSS (Hank’s Balanced Salt Solution) without calcium and magnesium (Gibco, Thermo Fisher Scientific). Five million cells were set aside as resting controls, while the remaining cells were centrifuged and resuspended in HBSS containing calcium and magnesium (Gibco, Thermo Fisher Scientific). For treatment, five million cells per condition were suspended in 1 mL HBSS and 1 µL of ionomycin stock dilutions (Sigma-Aldrich, dissolved in DMSO) were added at final concentrations of 0.01, 0.03, 0.1, 0.3, and 1 mM. Cells were incubated at 37°C for 30 minutes with gentle shaking. Following treatment, cells were lysed using 2% SDS in 40 mM Tris-HCl (pH 7.6) supplemented with 1X protease inhibitor cocktail (S8820, Sigma-Aldrich). Lysates were heat-inactivated at 90°C for 10 minutes, then acidified to 1% TFA for DNA hydrolysis and neutralized with 3M Tris buffer. Protein concentrations were determined using the BCA assay according to the manufacturer’s protocol. Twenty micrograms of protein from each sample was used for western blot analysis.

### Western blotting

Lyophilized samples (20–30 µg) were reconstituted in 30 µL of NuPAGE LDS sample buffer (Invitrogen) supplemented with 25 mM DTT. Samples were heated at 90°C for 10 minutes before loading onto NuPAGE 4–12% Bis-Tris gels (10- or 12-well, Invitrogen). PageRuler Plus Prestained Protein Ladder (Thermo Fisher Scientific) was used as a molecular weight marker, and NuPAGE MOPS buffer (Invitrogen) as the running buffer. Electrophoresis was carried out at 200 V for 45 minutes using an XCell SureLock Mini Cell chamber (Invitrogen). Proteins were transferred onto a 0.45 µm PVDF membrane (Merck Millipore) using NuPAGE transfer buffer (Invitrogen) and the XCell II blot module (Invitrogen) at 30 V for 1 hour. Membranes were stained with Ponceau S (Sigma-Aldrich) for 10 minutes to confirm equal loading and then destained with ultrapure water. Prior to antibody probing, membranes were washed three times with TBST and blocked with 2% BSA in TBST for 1 hour at room temperature. Membranes were incubated overnight at 4°C with primary antibodies under gentle shaking using 1) Custom antibodies: 12A2 and 21F10 (unpurified supernatant, 1:10); 2) Commercial antibodies: 07-377 (EMD Millipore, 1:1,000), MABN328 (EMD Millipore, 1:500), and ab100932 (Abcam, 1:500). Following three TBST washes, membranes were incubated with secondary antibodies (LI-COR, Inc.) for 30 minutes at room temperature: IRDye 800CW-coupled goat anti-rat IgG (926-32219); IRDye 800CW-coupled goat anti-rabbit IgG (926-32211); IRDye 680RD-coupled goat anti-mouse IgM (925-68180). All secondary antibodies were diluted 1:10,000 in TBST. After thorough washing with TBS, fluorescence detection was performed using a Li-Cor Odyssey Scanner (LI-COR, Inc.).

### Mass spectrometry analysis

#### Sample preparation

To confirm modifications, 100 µg of control, citrullinated, and homocitrullinated HeLa lysates were reconstituted in digestion buffer (100 mM HEPES, pH 8.5, 10 mM TCEP, 55 mM CAA) and incubated at 37°C for 1 hour with shaking for protein reduction and alkylation. Trypsin was added (1:50 enzyme:protein ratio), and samples were digested overnight at 37°C. Digests were acidified with 1% of formic acid and desalted using Sep-Pak C18 SPE cartridges (Waters). Peptides were eluted, lyophilized, and stored at −20°C until MS analysis.

#### Data acquisition

Samples were reconstituted in 10 µL of 0.1% FA and analyzed via LC-MS/MS on a Vanquish Neo LC system coupled to an Orbitrap Exploris 480 mass spectrometer (Thermo Fisher Scientific). A 5 µL injection (ca. 25 μg peptide) was loaded at 100 µL/min onto an Acclaim PepMap RSLC C18 column (Thermo Fisher Scientific), separated using a 60-minute gradient (mobile phases A: 0.1% FA, 3% DMSO in H₂O; B: 0.1% FA, 3% DMSO in ACN) at 50 µL/min, and ionized at 3500 V. The MS was operated in data-dependent acquisition mode with a 1.2 s cycle time, MS1 resolution of 60,000 (m/z 360–1300), and MS2 resolution of 15,000, using HCD fragmentation (28% collision energy) and dynamic exclusion (30 s, 10 ppm tolerance).

#### Database search and data analysis

Raw data were analyzed using FragPipe (v21.1) (MSFragger v4.0) against a human canonical protein database (Uniprot, 20,409 entries, downloaded 20.11.2023). Trypsin was set as the digestion enzyme (up to 3 missed cleavages, peptide length 7–50 amino acids, mass range 500–5000 Da). Fixed modification: carbamidomethylation (C); variable modifications: oxidation (M), acetylation (N-term), citrullination (R, +0.9840 Da), deamidation (N/Q, +0.9840 Da), and carbamylation (K, +43.0058 Da), allowing up to three modifications per peptide. Rescoring using MSBooster was disabled. All other search parameters were kept at default values. Modification levels were assessed by comparing peptide numbers and intensities. In the citrullinated sample, 21.5% of R-containing peptides and 9.1% of total peptides were modified (11.5% and 3.8% of intensity, respectively). The homocitrullinated sample showed near-complete lysine conversion, with 97.3% of K-containing peptides and 52.8% of total peptides modified (96.1% and 56.1% intensity-based). In contrast, modifications were detected in <0.1% of peptides in the control sample, within the 1% false discovery rate at PSM level.

### Motif Analysis

Potential substrate motifs around citrullination and homocitrullination sites were analyzed using pLOGO^23^. Sequences spanning five residues upstream and downstream of citrullination (R) or homocitrullination (K) sites were extracted and compared to background frequencies in the human proteomes. Statistically significant motifs (*p* = 0.05, Bonferroni corrected) were visualized.

## Results

### Generation of a peptide pool for immunization

Choosing appropriate immunogens is the key to generating successful antibodies for specific applications. Traditionally, three types of immunogens have been used to produce anti-pan citrullination antibodies: (1) single citrullinated peptide, (2) Single amino acid L-citrulline conjugated to carrier proteins like KLH or BSA, and (3) long chains of 10 consecutive citrulline^13^ (**Table 2**). These approaches share a common limitation: they do not account for the sequence context surrounding the modified site. Since antibodies typically recognize epitopes of 6–12 amino acids in length^24,25^, those generated using single peptide, protein, or non-contextual citrulline modifications may lack the broad specificity necessary to detect citrullinated sites with diverse adjacent residues.

Based on our prior identification of PAD2- and PAD4-specific motifs in human tissue proteomes^9^, we hypothesized that immunogens incorporating common motifs of citrullination sites could enhance specificity and reduce cross-reactivity with homocitrullination. To test this, we first synthesized a pool of diverse 10-residue peptides, incorporating 4 to 6 different amino acids per position (**Table 3**). These amino acids were selected based on their statistically significant enrichment (*p* < 0.05; Bonferroni corrected) at corresponding positions in citrullination motifs compared to human proteome. The percentages of amino acids at each position were weighted according to their observed frequencies, resulting in a pool comprising over 490,000 unique peptide combinations. This immunogen pool captures the most prevalent citrullination motifs in the human proteome, providing a robust foundation for generating motif-specific antibodies.

### Selection of specific monoclonal antibodies against a pool of citrullinated peptides

After immunizing rats with the citrullinated peptide pool, spleen cells from immunized animals were fused with a myeloma cell line to generate hybridoma clones for antibody production. Initial screening was conducted using bead-based flow cytometry (**Figure 1A**). In this assay, two pools of biotinylated peptides—either containing citrulline or arginine—were coupled to streptavidin beads conjugated with distinct fluorophores in the same reaction. Among the screened supernatants from 48x 96-well plates, two (12A2 and 21F10) demonstrated specific binding to citrullinated peptides while showing no interaction with their arginine counterparts, indicating high specificity.

To evaluate the ability of these clones to capture citrullinated peptides, a capture ELISA assay was performed. Plates were coated with subclass-specific anti-rat antibodies to immobilize the secreted antibodies in the hybridoma supernatants. Biotinylated citrullinated peptides were then added, and binding was detected using HRP-labeled streptavidin. Both clones exhibited significant binding to citrullinated peptides, confirming its capability to capture epitopes within the immunogen pool (not shown).

After limiting dilution cloning and expansion, the binding affinity of the two clones was further tested using a dilution series of biotinylated peptides containing either citrulline or arginine residues. These peptides were immobilized on avidin-coated plates and incubated with hybridoma supernatants. As shown in **Figure 1B**, both clones demonstrated strong specificity for citrullinated peptides, with negligible binding to arginine peptides. Notably, clone 21F10 displayed higher signal compared to clone 12A2, indicating its higher affinity.

### Reactivity of custom monoclonal antibodies and commercial antibodies toward citrullination and homocitrullination in western blotting

Given the specificity of 12A2 and 21F10 monoclonal antibodies toward citrullinated peptides, we further evaluated their ability to detect a broad range of citrullinated proteins in human proteomes. Proteins extracted from the HeLa cervical cancer cell line were treated overnight with recombinant PAD4 enzyme in the presence of calcium to induce citrullination, or with a high concentration of cyanate to induce homocitrullination. The presence of citrullinated and homocitrullinated proteins were confirmed by mass spectrometry (**Supplementary Table S1**). Notably, the identified citrullination sites aligned with the motif used in the immunogen design, whereas homocitrullination sites did not exhibit any specific motif preference (**Supplementary Figure S1A**). Western blotting was then performed to assess antibody reactivity toward these modifications. For comparison, three commercially available anti-pan citrullination antibodies—07-377, ab100932, and MABN328 (F95)—were included, as their reactivity to citrullinated and homocitrullinated antigens have been well-characterized^14,26^.

Both 12A2 and 21F10 specifically detected citrullinated proteins in PAD4-treated HeLa lysates (**Figure 2A; Supplementary Figure S2A**), with 21F10 displaying a particularly strong and specific signal and no detectable reactivity in negative controls. While 12A2 showed minimal background signal in the negative control and a faint band in the homocitrullinated sample around 60 kDa, both clones displayed markedly lower signal in homocitrullinated samples compared to citrullinated ones, demonstrating great specificity for citrullination.

**Figure 2.**
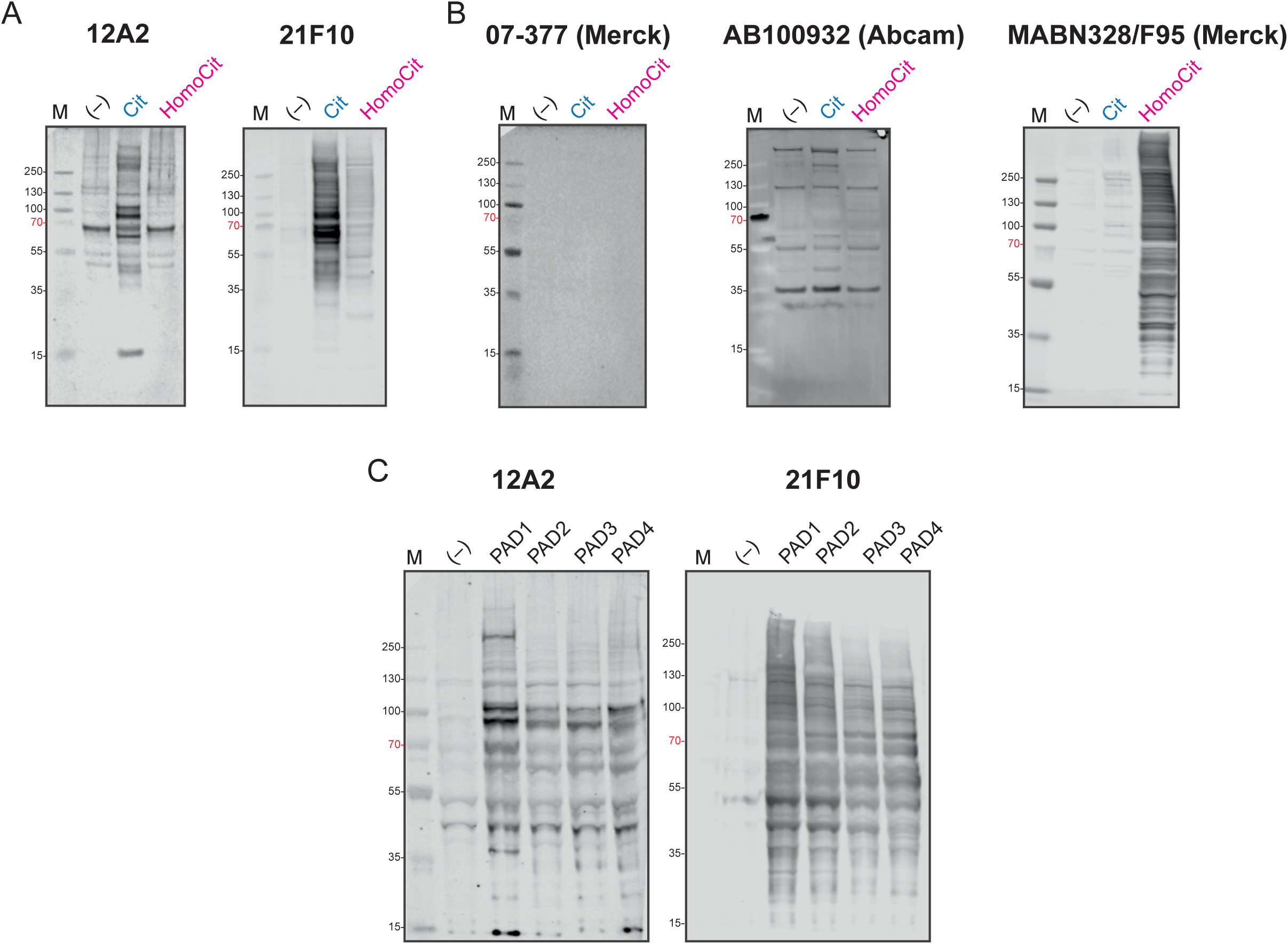
Reactivity of custom and commercial anti-pan citrullination antibodies in western blot analysis of in vitro citrullinated (Cit) and homocitrullinated (HomoCit) HeLa cell lysates. (A) Reactivity of motif-specific monoclonal antibodies 12A2 and 21F10 against in vitro citrullinated or homocitrullinated HeLa proteomes. (-), untreated HeLa lysate. (B) Reactivity of three commercially available anti-pan citrullination antibodies (07-377, ab100932, and MABN328 (F95)) against citrullinated and homocitrullinated HeLa lysates. (-), untreated HeLa lysate. (C) Reactivity of motif-specific monoclonal antibodies 12A2 and 21F10 toward HeLa proteomes treated with different PAD isozymes (PAD1, PAD2, PAD3, PAD4). (-), untreated HeLa lysate.

In contrast, the commercial antibodies exhibited varying performance (**Figure 2B; Supplementary Figure S2B**). The 07-377 antibody failed to detect any modified proteins, suggesting low sensitivity. The ab100932 antibody produced nonspecific signals even in negative control samples, aligning with the manufacturer’s recommendation that it can only be used exclusively for ELISA and immunocytochemistry (**Table 2**). MABN328 (F95) showed strong cross-reactivity toward homocitrullination. When probing citrullination and homocitrullination at comparable levels, F95 displayed even greater reactivity toward homocitrullinated proteins (**Figure 2B**). This observation is consistent with prior reports, where Verheul *et al.*^14^ demonstrated F95’s cross-reactivity with homocitrullinated fibrinogen and albumin, while Chen *et al.*^26^ noted its nonspecific binding to unmodified BSA.

These results highlight the superior specificity of the motif-based monoclonal antibody 21F10 for detecting citrullinated proteins, with minimal cross-reactivity toward homocitrullination. Their high specificity makes these antibodies valuable tools for investigating global citrullination in complex biological samples.

### Specificity toward citrullinated proteins catalyzed by different PAD enzymes

Since the motifs represented in our immunogens were primarily derived from tissues expressing PAD2 and PAD4^9^, we sought to determine whether the reactivity of the 12A2 and 21F10 clones is broadly applicable to citrullination catalyzed by various PAD enzymes. To assess this, HeLa lysates were incubated with recombinant PAD1, PAD2, PAD3, or PAD4 in the presence of calcium, and the resulting citrullinated proteomes were analyzed via western blotting.

Both clones exhibited robust reactivity across all PAD-treated samples, with slightly stronger signals observed in PAD1-treated proteomes (**Figure 2C; Supplementary Figure S2C**). However, there was no distinct difference in citrullination between samples treated with different PAD enzymes whereas the citrullinated proteins recognized by the two clones were different.

These findings suggest that 12A2 and 21F10 broadly recognize PAD-catalyzed citrullination, with differences in motif recognition, making them valuable for studying citrullination in diverse biological contexts where different PAD isozymes are active.

### Detection of global changes in citrullination during neutrophil activation

The above analyses demonstrate that 12A2 and 21F10 recognize a wide array of citrullinated substrates, with 21F10 showing a higher sensitivity. However, its sensitivity in detecting citrullination in biological samples remains unclear due to the artificially high levels of citrullination in *in vitro* experiments (∼9-10%). To evaluate the ability of 21F10 to detect (semi-)quantitative changes in global citrullination, we isolated human neutrophils and treated them with the calcium ionophore ionomycin in a dose-dependent manner. This treatment is known to activate PAD enzymes, elevate citrullination levels, and induce neutrophil extracellular trap (NET) formation^27^.

We detected dose-dependent increase of signal in global citrullination levels by 21F10 (**Figure 3**). These results indicate that 21F10 can effectively recognize changes in pan-citrullination associated with PAD enzyme activation in biological samples in a quantitative manner, showing its potential utility for studying dynamic citrullination processes.

**Figure 3.**
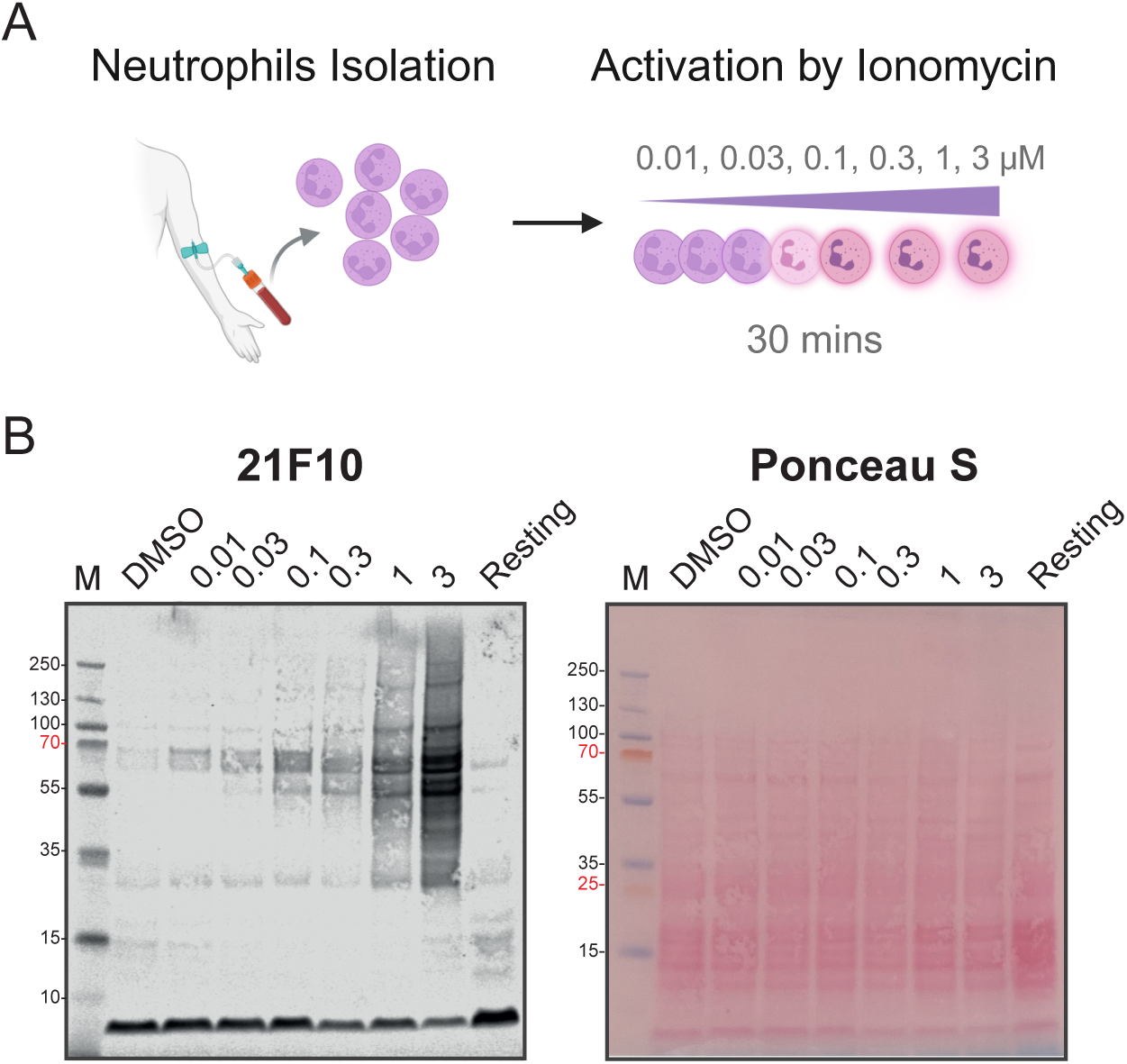
Detection of global citrullination changes during neutrophil activation. (A) Schematic representation of neutrophil activation by ionomycin treatment. (B) Reactivity of motif-specific monoclonal antibodies 21F10 against citrullinated proteins in ionomycin-activated neutrophil proteomes. Left panel, citrullination signals detected by western blot; Right panel, Ponceau S staining confirming equal protein loading across samples.

## Discussion

Affinity-based detection of target proteins is indispensable in biological research. However, antibody performance varies greatly depending on the target protein and its modifications^28^. Detecting global level of PTMs via antibodies is particularly challenging, as it requires recognizing a shared epitope independent of the surrounding sequence context. The ubiquitin remnant motif is one successful example, where a motif-specific antibody has been developed to detect ubiquitinated peptides in a sequence-independent manner^19^.

Here, we introduce a new class of anti-pan citrullination antibodies based on motifs preferentially modified by PAD enzymes in the human proteome. We assessed their specificity and cross-reactivity within complex biological samples using western blotting. Through comparative analysis with three commercially available anti-citrullination antibodies, we demonstrated that our custom monoclonal antibodies—particularly clone 21F10—exhibit superior specificity and sensitivity toward citrullinated proteins while displaying minimal cross-reactivity with homocitrullinated proteins. Furthermore, we validated the ability of 21F10 to detect dynamic changes in protein citrullination during neutrophil activation, underscoring its potential as a valuable tool for studying citrullination in biological systems.

Despite their promising performance, several aspects warrant further investigation. First, the observed differences in signal patterns between 12A2 and 21F10 suggest that these clones recognize distinct citrullinated motifs. A combinatorial approach using multiple monoclonal antibodies may therefore enhance detection coverage and further improve overall sensitivity. Second, while we evaluated widely used commercial antibodies with well-characterized performance in ELISA (07-377, ab100932, MABN328) and immunoblotting (MABN328)^14^, we did not cover all reported anti-pan citrullination antibodies. Some lack validation data, while others are not commercially available^26^. This reflects a broader challenge in the field: the absence of standardized validation protocols for anti-pan citrullination antibodies and the limited accessibility of well-characterized reagents. Establishing reference standards, such as *in vitro* citrullinated lysates or PAD-activated cell models and ensuring the availability of validated antibodies through reputable research centers or vendors would be instrumental for enhancing reproducibility and broadening accessibility across studies.

While our findings demonstrate the feasibility of a motif-based strategy for generating highly specific and sensitive anti-pan citrullination antibodies, certain limitations must be considered. First, although our peptide pool design incorporated over 490,000 motif combinations to comprehensively represent citrullination sites, rare motif variants may not be fully captured, potentially limiting complete proteome coverage. Second, although our custom antibodies showed minimal cross-reactivity with homocitrullinated proteins, structural similarities between citrullinated and homocitrullinated motifs could still lead to occasional off-target binding. In biological samples with exceptionally high levels of homocitrullination, the resulting signal could obscure citrullination-specific detection. Although such scenarios may be uncommon, integrating mass spectrometry-based approaches to quantify and distinguish between these modifications would help to address this limitation. Notably, mass spectrometry remains indispensable for precisely characterizing citrullination sites and differentiating them from homocitrullination. Future studies combining high-quality antibodies with advanced proteomic techniques will provide a more comprehensive understanding of citrullination in both physiological and pathological contexts.

We believe that the continued development of reliable anti-pan citrullination antibodies has the potential to advance fundamental research on citrullination, facilitate biomarker discovery, and improve disease diagnostics. By enabling more accurate detection of citrullinated proteins, these tools may contribute to the identification of novel disease mechanisms and the development of targeted therapeutic strategies.

## Resource availability

### Materials availability

Both clones, 12A2 (RRID: AB_3677762) and 21F10 (RRID: AB_3677763), generated in this study are available on request, but we may require a payment and/or a completed materials transfer agreement if there is potential for commercial application. Please contact the lead contact Chien-Yun Lee (.) and Regina Feederle.

### Data availability

The mass spectrometry proteomics data have been deposited to the ProteomeXchange Consortium via the PRIDE^29^ partner repository with the dataset identifier PXD061971.

## Acknowledgments

The authors would like to express their gratitude to all members of the Lee Lab, the Bavarian Center for Biomolecular Mass Spectrometry (BayBioMS), and the Chair of Proteomics and Bioanalytics for their valuable assistance and insightful discussions. This work was funded by the Federal Ministry of Education and Research (FKZ031L0215; YIG-SysNS). The Exploris 480 mass spectrometer was funded in part by the German Research Foundation (DFG-INST 36/171- 1 FUGG). The graphical abstract and illustrations were created with BioRender.com.

## Author contributions

**Sophia Laposchan:** Data curation, Investigation, Methodology, Validation, Writing – original draft, Writing – review & editing; **Erik Riedel:** Data curation, Investigation, Validation; Andrew Flatley: Investigation, Methodology**; Regina Feederle**: Conceptualization, Investigation, Methodology, Project administration, Writing – review & editing; **Chien-Yun Lee:** Conceptualization, Investigation, Methodology, Funding acquisition, Project administration, Supervision, Writing – original draft, Writing – review & editing.

---

**Supplemental Figure 1.**
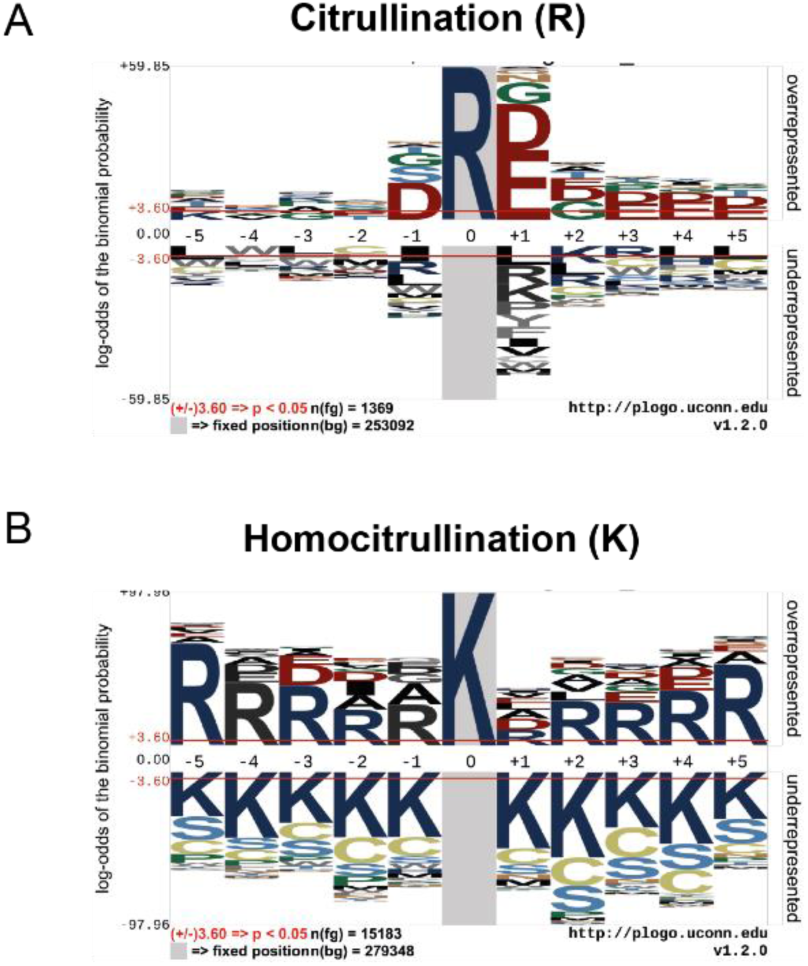
Motif analysis of citrullinated and homocitrullinated peptides identified in HeLa lysates. (A) Sequence motif analysis of citrullination sites. (B) Sequence motif analysis of homocitrullination sites.

**Supplemental Figure 2.**
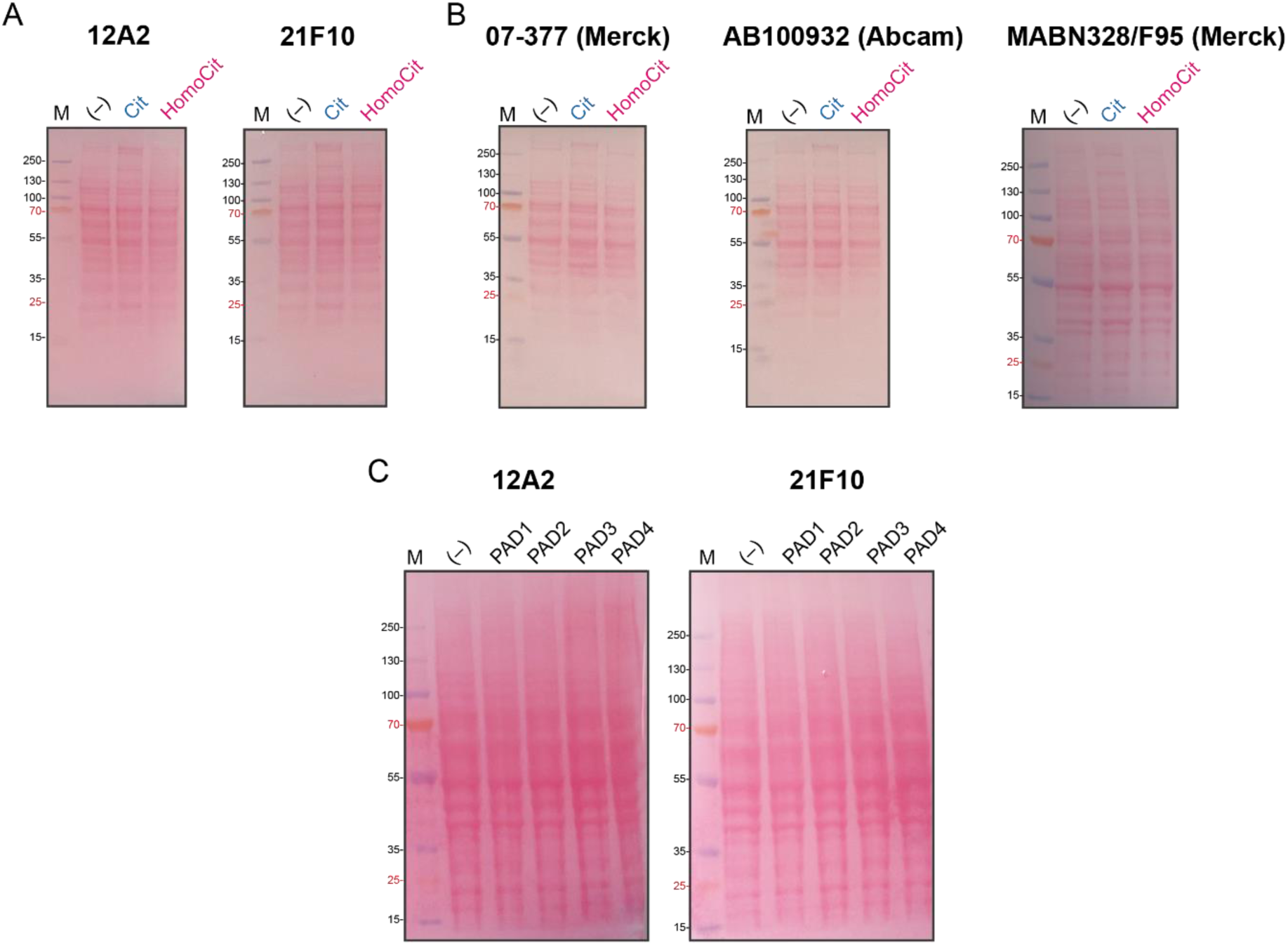
Ponceau S staining. The plot is corresponding to the blots shown in Figure 2, confirming equal protein loading.

